# Decoding instrumental lever pressing from prefrontal, accumbens, and hippocampal local field potentials: conservation across sex, task, and dopamine depletion

**DOI:** 10.64898/2026.09.16.751848

**Authors:** Abhijith Mankili, Alev Ecevitoglu, Gayle A. Edelstein, Naixin Ren, Renee A. Rotolo, James J. Chrobak, John D. Salamone, Ian H. Stevenson

## Abstract

During operant behavior, oscillatory activity is coordinated across multiple brain regions and can change under different task conditions. In this study, we use a decoding approach to characterize these patterns and predict lever pressing behavior in rats from local field potentials (LFP) recorded bilaterally in the hippocampus, nucleus accumbens, and prefrontal cortex. We extract LFP features for band power and peak frequency and predict behavior across animals and conditions with a Poisson Generalized Linear Model (GLM). We use data from both male and female rats performing either fixed-ratio (FR40) or progressive (PROG) operant lever-pressing tasks, under vehicle (VEH) or tetrabenazine (TBZ), a vesicular monoamine transport (VMAT-2) inhibitor that depletes dopamine and induces depressive-like motivation dysfunction. We find that we can accurately predict lever pressing from multi-region LFP within-animals on a timescale of seconds with >30% variance explained. Although within-animal and within-condition predictions are highest, we also find, interestingly, that these decoders generalize across animals and across drug, sex, and task conditions, suggesting a stable association between LFP and behavior. We then evaluate whether LFP features can be used to decode long-term task variables, such as the number of presses since or until the next reinforcer. We find that LFP features can predict these long-term task variables, but that decoding relies on different LFP features for prediction than those used for immediate lever pressing. Altogether these results suggest that there are robust distributed patterns of LFP associated with lever-pressing behavior under multiple drug, task, and sex conditions.

**Significance Statement:** Although patterns of neural activity related to many observable behaviors, such as sleep and locomotion, have been well described, identifying stable neural patterns related to other states, such as motivation and effort, is an ongoing challenge. Here we use a decoding approach and aim to predict effort-based lever pressing from neural activity for a wide range of conditions, with rats of different sexes performing different tasks with and without a drug, tetrabenazine. We find that there is a stable pattern of neural features that allows us to robustly predict lever-pressing across these many conditions. This pattern may serve as a general indicator of effortful behavior, and deviations from the pattern could potentially indicate motivational dysfunction.

## Introduction

Brain-wide patterns of oscillatory activity, referred to as local field potentials (LFP), provide important indicators of behavior and neurological disorders (Ward, 2003; Buzsáki et al., 2012; Weiss et al., 2023; Van Bree et al., 2025). Specific LFP features have been robustly associated with behaviors in many systems, such as the alpha rhythm associated with attention and arousal (Klimesch, 2012; Jensen and Bonnefond, 2026) or the theta rhythm associated with locomotion (Vanderwolf, 1969; Whishaw and Vanderwolf, 1973; Buzsáki, 2002; Long et al., 2014). However, the LFP features associated with operant behavior and motivational states are less well established. Previous work has found that the power of both the delta and theta rhythms, particularly in the hippocampus, is associated with lever pressing in rats (Gruber et al., 2009; Ecevitoglu et al., 2026a), but whether these trends are robust across conditions and animals has not been fully established. Here we characterize LFP activity in three interconnected brain regions: the hippocampus (HpC), prefrontal cortex (PFC), and nucleus accumbens (NAc) network, during two goal-directed operant lever-pressing tasks. We use a decoding approach (Mathis et al., 2024), where we aim to predict lever pressing using multiple LFP features from these three brain areas. Since LFP-behavior associations in many systems differ across tasks or pharmacological manipulations, we aim to identify which LFP features are conserved across contexts and to determine whether a common, multi-region pattern of LFP features allows behavior to be predicted across contexts.

The HpC, PFC, and NAc play key roles in goal-directed behavior (Mogenson et al., 1980; Goto and Grace, 2008; Gruber et al., 2009), and PFC (Hauber and Sommer, 2009) and NAc (Salamone et al., 2003, 2007; Mai and Hauber, 2012), in particular, have been linked to motivational processes. These regions are interconnected, with PFC and HpC providing glutamatergic input to NAc (Britt et al., 2012), and all three areas are also primary targets for dopamine pathways in the brain (Björklund and Dunnett, 2007; Sesack and Grace, 2010). Dopaminergic signaling is essential for the execution of motivated behavior (Salamone et al., 2007, 2018; Salamone and Correa, 2024), and pharmacological manipulations that decrease dopamine levels cause motivational deficits (Salamone et al., 2016). One such drug is tetrabenazine (TBZ), a vesicular monoamine transporter 2 (VMAT-2) inhibitor (Scherman et al., 1988) that acts as a dopamine depleting agent, particularly in the striatum, leading to depressive symptoms in animals including humans (Chen et al., 2012; Nunes et al., 2013; Salamone and Correa, 2024). Previous work found that TBZ administration produced a low-effort bias in rats, reducing lever-pressing in rats tested on operant ratio schedules (Nunes et al., 2013; Randall et al., 2014; Ren et al., 2022). Interestingly, the effective dosage of TBZ appears to differ between male and female rats, with females requiring a higher dosage to induce comparable behavioral effects (Ecevitoglu et al., 2024a). Similarly, training schedules (Johnson et al., 2022) and the specific behavioral dynamics of different task ratios (Aberman and Salamone, 1999; Hamill et al., 1999), may result in differential sensitivity to dopaminergic manipulations. These effects raise the possibility that there are task, drug, and sex-specific associations between behavior and neural activity in the HpC, PFC, and NAc. We, thus, aim to identify which patterns of neural activity are conserved and which vary under these different conditions.

Identifying the specific features of multi-region LFP that are associated with behavior is a challenging statistical problem (Einevoll et al., 2013; Pesaran et al., 2018; Van Bree et al., 2025). Here we focus on time-varying power and peak frequency within a defined set of frequency bands: delta, theta, beta, low-gamma, and high-gamma. LFP activity within these bands reflects distinct neural processes (Buzsáki et al., 2012) and both the power and frequency within these bands has been previously associated with specific behavioral functions (Besosa et al., 2026). However, since these features co-vary, their independent associations with behavior may be misleading.

We instead consider a model-based approach, where we aim to predict or decode behavior from all simultaneously observed LFP features (Jackson and Hall, 2017). Decoding approaches have been used to identify how well behavior can be predicted from neural activity and which features or brain areas are most informative (deCharms and Zador, 2000; Kriegeskorte and Douglas, 2019). These models can be used to determine whether the LFP-behavior relationships in one condition, such as a specific animal of a specific sex performing a specific task with a specific drug, generalize to different settings.

Here we decode multi-area LFP features to predict operant lever pressing behavior in rats. We measure both within and across animal prediction accuracy and find that press rate can be predicted from band power and peak frequency features in the HpC, NAc, and PFC in a wide range of contexts. Across animals, we find a conserved pattern of predictors that weighs HpC theta activity and NAc gamma activity most heavily and seems to occur consistently across conditions.

## Materials and Methods

Here male and female rats were trained to perform an operant lever pressing task. After initial training, animals were implanted bilaterally with 50-micron tungsten microwire electrodes in prefrontal cortex, nucleus accumbens, and dorsal hippocampus. Animals were then retrained and subsequently recorded performing the operant task under both vehicle and tetrabenazine conditions.

All experimental procedures were approved by the University of Connecticut Animal Care and Use Committee and in accordance with NIH and the American Veterinary Medical Association guidelines.

### Experimental design

#### Subjects

A total of 26 male and 25 female Sprague Dawley rats (Envigo, Indianapolis, IN) age ∼3-6 months were used. The animals were housed in a vivarium maintained at 23 ºC and on a 12-h light/dark cycle. The animals were habituated to the colony for a week followed by food-restriction to maintain 85% of their free-feeding body weight initially, followed by adherence to a growth curve that allowed modest growth across the experiments. The animals then underwent operant training, by the end of which the male rats on average weighed ∼305g and female rats on average weighed ∼265g. Following surgery and retraining, average weights at the time of postsurgical testing was ∼340g for males and ∼278g for females. Rats were single-housed during surgical recovery and for the remaining duration of the experiment. Environmental enrichment (e.g., toys or shelters) was not provided following electrode implantation to avoid any interference with electrodes or surgical site.

#### Training

Animals were trained to perform one of two tasks: a fixed-ratio 40 (FR40) task and a progressive ratio (PROG) task. Following previous training procedures (Ecevitoglu et al., 2026a), rats underwent behavioral sessions (30 min/session, 1 session/day, 5 days/week) in 28×23×23 cm operant chambers (Med Associates, Fairfax, VT) and were trained to lever press for high-carbohydrate 45 mg pellets (Bio-Serv, Flemington, NJ). Training began with magazine habituation for 3 days, followed by an FR1 schedule for one week. After acquiring FR1, FR40 animals were shifted to an FR20 schedule for one week and then trained on the FR40 schedule for five additional weeks. PROG animals, after acquiring FR1 were trained on the PROG schedule where the ratio started at FR1 and was increased by one additional response for every 15 deliveries of the reinforcer. PROG training was continued for 9 weeks during which the session ended early if the animal did not complete a ratio within 2 minutes.

#### Pharmacological agent

TBZ was acquired from Tocris Bioscience (Ellisville, MO) and was prepared by dissolving in DMSO (20%) and 0.9% saline (80%) followed by titration with 1.0 N HCl (pH of 4.0–4.5), The drugs were administered intra-peritoneally (IP) 120 minutes before testing. VEH included DMSO, saline and HCl to be used as control. TBZ was administered at 1.0mg/kg dosage for males and 2.0mg/kg for females.

#### Surgery

Animals were implanted with 16 custom-made 50-micron tungsten microwire electrodes following the coordinates and procedures described in (Ecevitoglu et al., 2026a). Briefly, holes were drilled according to the following coordinates from bregma: PFC: AP (+) 3.8 mm, ML (±) 1.4 mm; NAc: AP (+) 1.0 mm, ML (±) 1.4 mm; dorsal HpC: AP (−) 3.5 mm, ML (±) 1.4 mm. Electrodes were pre-cut and lowered to DV (−) 3.2 mm (PFC), DV (−) 7.2 mm (NAc), and DV (−) 4.2 mm (HpC), with the NAc used as the anchor point to determine DV placement. Coordinates and depths were identical across sexes. After one week of recovery, animals were retrained.

#### Recording

Electrophysiological data were collected while the animals performed the effort-based lever pressing task using the digital Lynx SX Electrophysiology System (Neuralynx) software with a tethered cable used for capturing wide-band activity at a sampling rate of 2000 Hz. Data was low-pass filtered to isolate the LFP band 0.1-500Hz (FIR filter with 256 taps).

At the end of initial training (∼8 weeks, 1 session/day), animals underwent two pre-surgical behavioral recording sessions, during which lever-pressing behavior was recorded while performing their assigned task (FR40 or PROG) for 30 min under either TBZ or VEH, counterbalanced across animals in a within-subject design (one drug condition tested per week). Animals then underwent electrode implantation surgery, followed by one week of recovery and retraining on their assigned schedule. Following retraining, animals were habituated to the recording procedure for 2-3 days and then underwent two additional weeks of behavioral recording sessions under TBZ and VEH, following the same procedure as the pre-surgical sessions. Lever pressing under VEH and TBZ was similar for the pre-surgical and post-surgical behavioral sessions (Fig S1). At the end of the study, brains were harvested for histological assessment.

#### Histological assessment

Following completion of the study, rats were euthanized and perfused following the procedure described in (Ecevitoglu et al., 2026a).Briefly, animals were euthanized using CO_2_ anesthesia and underwent trans-cardiac perfusion with 0.9% saline followed by 3.7% formaldehyde. Brains were extracted and sectioned into 50-μm slices using a vibratome (Leica, USA), then stained with cresyl violet (Nissl stain) to verify electrode placement. Electrode tip locations were confirmed using the Paxinos and Watson Rat Brain in Stereotaxic Coordinates atlas, referenced against the coordinates listed in surgery (Fig S2). Sectioning and staining protocols were identical across sexes.

### Statistical analysis

#### Data preprocessing

Recordings were analyzed using MATLAB (MathWorks Inc, Natick, MA) with the FieldTrip (Oostenveld et al., 2011) and Chronux (Bokil et al., 2010) toolboxes. LFP data was first passed through an artifact detection pipeline to exclude signal clipping, large movement, and other noise artifacts. Data surrounding artifacts was also excluded by smoothing the artifact locations using a Gaussian moving window (3s SD) and removing data >10% of max of the smoothed signal. We visually assessed the artifact-removed data for quality and excluded channels with large fractions of artifacts, electrical disconnection, and severe 60Hz line noise. Some recording sessions had too many artifacts or too little pressing for reasonable decoding. Of the 102 total recordings (52 from male animals, 50 from female), we excluded 19 recordings from further analysis.

#### Predictors

We next extracted power and peak frequency features from the LFP to be used as predictors in our decoding model. We use a 25-order least-squares FIR lowpass filter to extract 0-4Hz (delta) and bandpass FIR filters to extract 6-12Hz (theta), 12-30Hz (beta), 30-58Hz (low-gamma), and 62-117Hz (high-gamma). Bandpass filters were designed using a Kaiser window with 1dB passband ripple and 60dB stopband attenuation. The stop band frequencies were spaced 1Hz above and below the respective upper and lower passband frequencies. We then estimate the power spectrum of each filtered signal in 2s non-overlapping bins to extract the power and peak frequency within each band. Peak frequency is based on the first conditional spectral moment of the time-frequency distribution. In total, we get 60 predictors corresponding to power and frequency features across all the 5 frequency bands for each of the 3 brain regions in both hemispheres.

#### Decoding

Predictors described above were z-scored for each animal and used in a Poisson generalized linear model (GLM) to predict the number of lever presses in 2s segments of the data (McCullagh and Nelder, 1989)

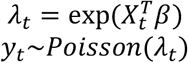

where *y*_*t*_ denotes the observed number of presses at time *t* and is predicted using a vector of LFP predictors *X*_*t*_ (including a constant predictor for the intercept) weighted by the vector of coefficients *β*. To fit coefficients, we maximize the penalized log-likelihood, and to prevent overfitting, we used L2-regularization and coarsely optimized the regularization penalty to maximize the performance on held-out data (penalty *η* = 1). We evaluated model performance within-animals using 10-fold cross-validation. After fitting the model coefficients to training data, we measure the McFadden’s pseudo-R^2^ on test data

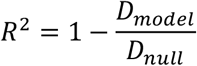

where *D* = −2[log(*L*_*model*_) − log(*L*_*saturated*_)] denotes the model deviance based on the log-likelihood log(*L*) ∝ ∑_*t*_ *y*_*t*_ log(*λ*_*t*_) − *λ*_*t*_. *D*_*null*_ corresponds to the deviance of the null model where *λ*_*t*_ is constant, and the deviance for both the full model and the null model are based on the difference between the log likelihood of that model and the log likelihood of the saturated model that uses *λ*_*t*_ = *y*_*t*_. Since the timeseries data has strong autocorrelations, we specifically used block cross-validation with a block size of 30s to reduce overestimating the performance from neighboring time points.

In the results above, each recording is optimized separately, resulting in unique fits for each animal and recoding. To evaluate the generalization of models across animals, we evaluate the performance of population models fit to multiple animals and then evaluate test performance to held-out recordings within and across groups. We consider population models fit to each combination of sex, task, and drug. We tested within-group generalization on held-out animals from the same combination of conditions, and across-group generalization on held-out animals that differ in one factor (sex, task, or drug). Note that these population models generally cannot predict any individual differences in overall pressing, since the held-out recordings are entirely missing from the training data. To evaluate the accuracy, we thus rescale the population predictions using

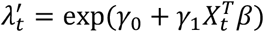

This secondary model uses the coefficients *β* of the population model to weigh the specific LFP features, but, additionally, adds an intercept *γ*_O_ and scaling *γ*_1_ to adapt the population predictions to each individual recording.

#### Long-term task variables

We also evaluated whether the LFP features can account for other aspects of the lever-pressing task besides the immediate press count. In particular, we calculated two variables that reflect where the animal is within each ratio: 1) number of lever presses executed since the previous reinforcer and 2) number of presses remaining until the next reinforcer. These variables represent cumulative measures of the amount of lever presses completed or remaining for the animal within each ratio, and, since they are still counts, we use the same GLM approach described above to assess whether they can be predicted from LFP features both within and across animals.

#### Coefficient vector geometry

To compare the similarity of coefficient vectors across conditions and animals we use cosine similarity (excluding the intercept). To identify patterns in the coefficients across all recording sessions we used nonmetric multi-dimensional scaling with cosine dissimilarity as the distance metric, again excluding the intercept. Here we use a 1-d embedding, with Kruskal’s normalized stress criterion.

## Results

To characterize the associations between neural activity and operant lever-pressing under a wide range of conditions, we record local field potentials (Fig 1A) during two operant tasks, from both male and female rats, after IP injections of the dopamine depleting agent tetrabenazine (TBZ) and the vehicle (VEH) control. Tasks included a fixed-ratio 40 task where 40 presses result in delivery of a food pellet (FR40) and a progressive ratio task where the number of lever presses required to get the pellet progressively increases throughout the behavioral session (PROG) During behavior, we recorded LFP simultaneously and bilaterally from three brain regions (Fig 1B): dorsal hippocampus (HpC), nucleus accumbens (NAc), and prefrontal cortex (PFC). We then analyze to what extent features of the LFP correlate with moment-by-moment lever pressing (Fig 1C).

**Figure 1.**
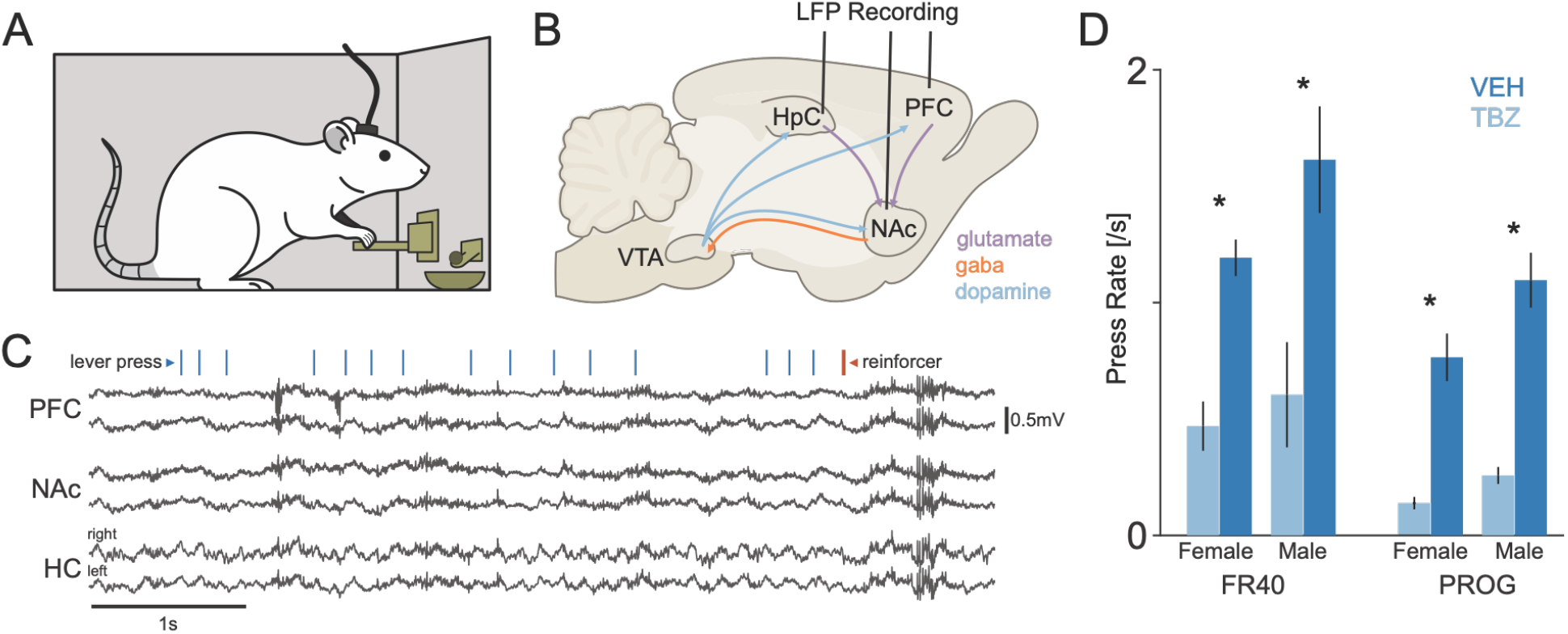
Task and recording framework. A) Multi-area Local Field Potential (LFP) signals were recorded bilaterally from the rat while the animals were engaged in an effort-based lever pressing task. B) Recordings targeted Hippocampus, Nucleus Accumbens and Prefrontal Cortex using chronically implanted single channel tungsten electrodes. C) Continuous LFP traces illustrate the temporal and spatial variability associated with behavior. Ticks indicate the timing of individual lever presses during ∼5s of behavior. Red tick indicates a lever press that resulted in the delivery of a pellet reinforcer. These example traces were recorded from a male rat performing FR40 task under vehicle condition. D) Average observed press rate for each experimental condition varying task, sex, and drug. Error bars denote standard error across recording sessions. Drug conditions include repeated measures within animals, and there are n=83 recordings in total. * Indicates p<0.05 (paired t-test).

First, to evaluate the relative impact of drug, task, and sex on overall behavior, we compare the press rate for all groups (Fig 1D). Previous work has found that TBZ results in substantial reductions in pressing in a wide range of conditions (Nunes et al., 2013; Yohn et al., 2016; Ren et al., 2022; Ecevitoglu et al., 2024b, 2025, 2026b), but that female rats are less sensitive to TBZ than male rats (Ecevitoglu et al., 2024a). To standardize the behavioral effects, we thus use different dosages for the male rats (1.0 mg/kg) and female rats (2.0 mg/kg) here. We analyze average lever press rates per session for the n=83 recording sessions here with a linear mixed effects model with random intercepts for individual animals. Task and sex comparisons are between animals, while the drug effect is measured within animals with counter-balanced recordings under VEH and TBZ. Consistent with previous studies, we find that there are substantial reductions in pressing rate under TBZ compared to VEH (F(1,75)=30.5, p<10^-6^) with an estimated marginal mean of 0.40 presses/sec for TBZ and 1.13 presses/sec for VEH. Pressing rate during PROG is also generally lower compared to FR40 (F(1,75)=5.5, p=0.02) with an estimated marginal mean of 0.57 presses/sec for FR40 and 0.22 presses/sec for PROG. However, we do not find statistically significant differences between male and female press counts for the data here (F(1,75)=0.67,p=0.42, 0.48 presses/sec female and 0.65 presses/sec male) or interactions between drug, task, and sex factors (p>0.05).

Simultaneously recorded LFP varies across brain areas and with lever pressing (Fig 2). Across recording conditions, we generally find that PFC and NAc have similar spectra, while HpC has a prominent peak in the theta band (6-12Hz). Additionally, both the power and peak frequency of the HpC theta band covary with press rate. This association suggests that lever pressing can likely be decoded from LFP features, and here we extract both power and peak frequency in five bands: delta (0-4Hz), theta (6-12Hz), beta (12-30Hz), low gamma (30-58Hz), and high gamma (62-117Hz). Each of these features is potentially associated with lever pressing behavior. Note, however, that there is substantial behavioral variability across task and drug conditions. PROG sessions typically show a gradual increase in press rate over the course of the session as the animal is required to press with increasing ratios before the reinforcer is delivered. TBZ sessions typically show substantially reduced pressing and more intermittent lever pressing over the course of the session.

**Figure 2.**
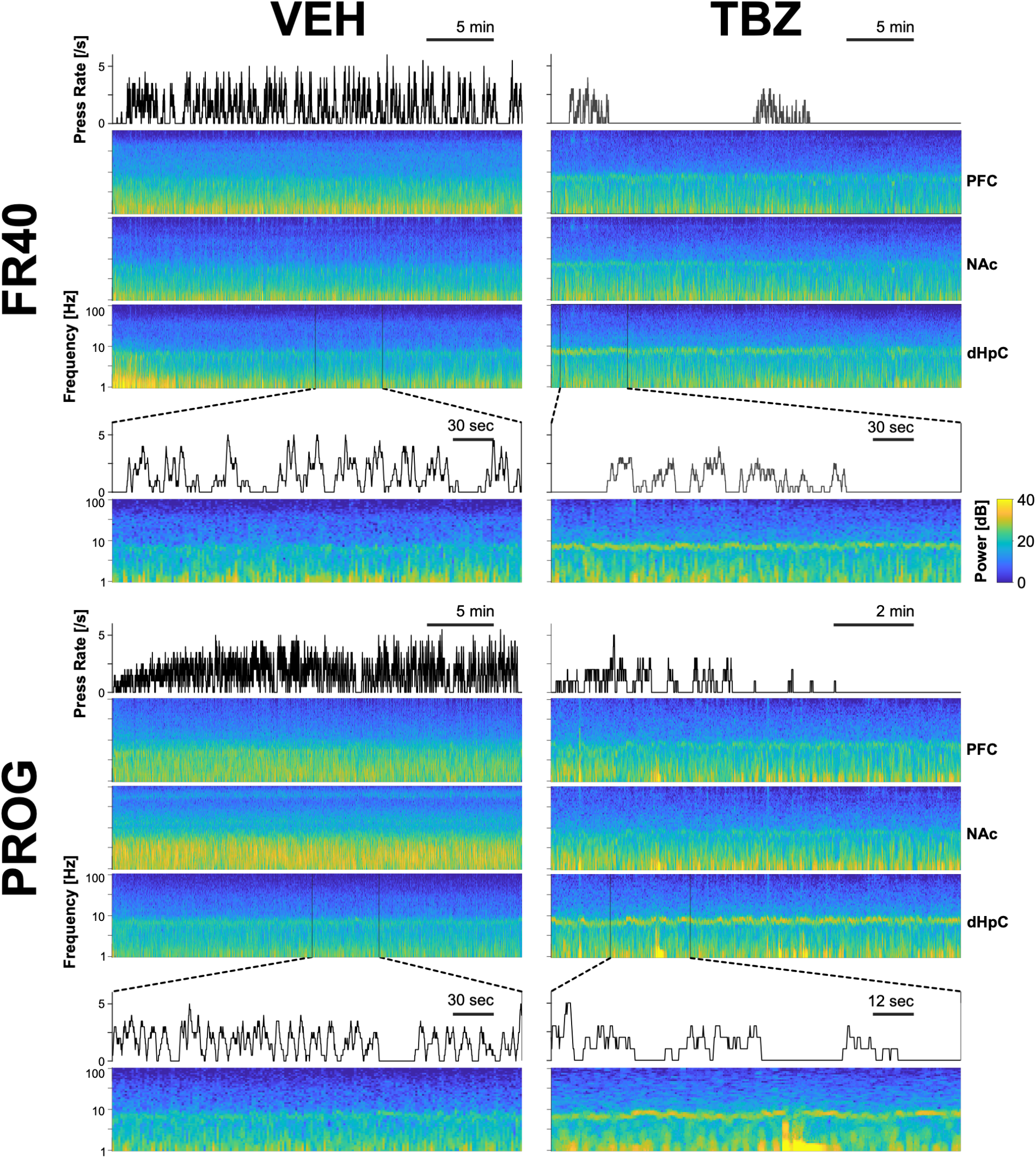
Lever pressing and spectro-temporal LFP patterns across tasks and drug conditions. Four example recordings are shown for the fixed-ratio 40 (FR40) task (top), progressive ratio (PROG) task (bottom), under vehicle (VEH, left) and tetrabenazine (TBZ, right) conditions. On the timescale of the ∼30min recording, the LFP spectrum is stable in prefrontal cortex, nucleus accumbens, and hippocampus. On the shorter timescale of the behavior, the spectrum of the LFP covaries with the rate of lever pressing. Most noticeably, the power in the delta (0-4Hz) and theta (6-12Hz) bands of the hippocampal channels and the peak frequency in the theta band tends to increase during periods of low lever pressing. The spectrogram and press rate shown here are both calculated in non-overlapping 2s bins. For reference, FR40 VEH and PROG TBZ data are from female animals, and FR40 TBZ and PROG VEH data are from male animals.

Here we use a decoding approach and predict press counts from neural signals within 2s windows. We used 60 different features – covariates such as band power and peak frequency – extracted from the LFP data of the animals as predictors in a Poisson-GLM to predict the local lever pressing rates of each individual animal (see Methods, Fig 3A).

**Figure 3.**
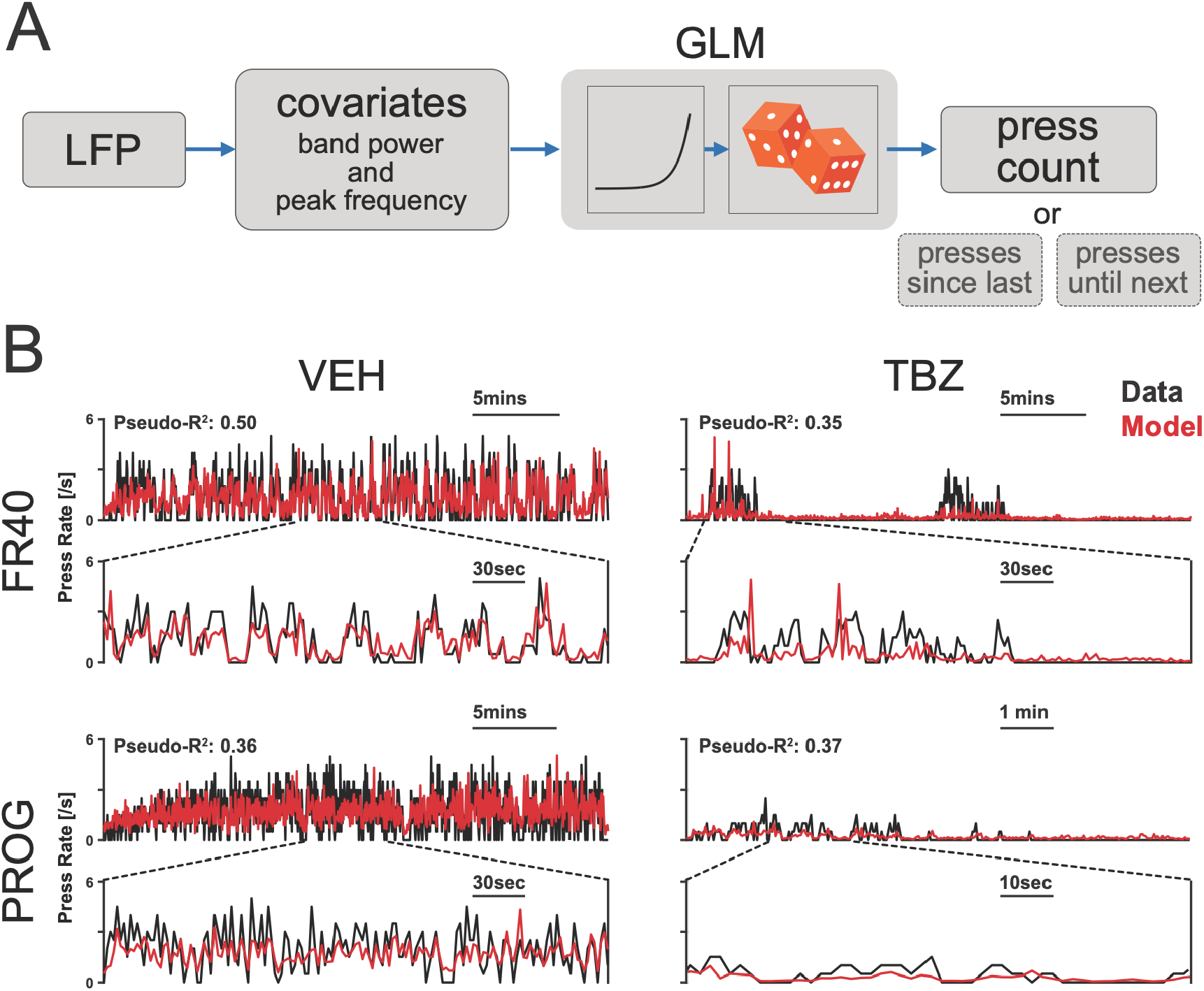
Decoding press rate from LFP features. A) Using frequency and amplitude features extracted from the LFP data, we use them as inputs of a Generalized Linear Model and predict the lever-pressing behavior of the animal, in-particular, the local press rate. We also use the model to predict two additional variables generated from the lever pressing timeseries data corresponding to long-term representation of the behavior: the number of presses since the last reinforcer and the number of presses until the next reinforcer. B) Example traces for observed (black) and predicted (red) lever pressing. Model traces and goodness-of-fit denote cross-validated predictions from the full model with all LFP predictors. These examples correspond to the data shown in Fig. 2.

We find that, in general, these LFP predictors can predict the moment-by-moment changes in press rate. To prevent overfitting, the model was regularized, and we measure accuracy using 10-fold block cross-validation with a block size of 30s. The overall within-animal psuedo-R^2^ using all predictors is 36.5±0.1% (mean±SE). Here we assume that the press count observations in 2s windows are well described by a Poisson distribution. However, the observed count distributions often have greater occurrence of 0 pressing than predicted by the Poisson model.

To evaluate the robustness of these LFP-based decoders we fit models with only subsets of features (Fig 4A). Although the model with all (60) features performs best, using subsets of features result in only modest drop in accuracy. Using only specific brain regions, we find that the LFP features from the hippocampal channels are most informative with a pseudo-R^2^ of 29.4±0.1%. Similarly, using only specific frequency bands, we find that the theta band features (across all regions) are most informative pseudo-R^2^ of 22.4±0.1%. Using power features (30) gives slightly better performance (29.1±0.1%) than using peak frequency features (24.3±0.1%), and using features from the right hemisphere gives slightly better performance (31.2±0.1%) than using features from the left hemisphere (29.4±0.1%). However, the most accurate model includes all features, suggesting that information from multiple regions, bands, hemispheres, and feature types is useful.

**Figure 4.**
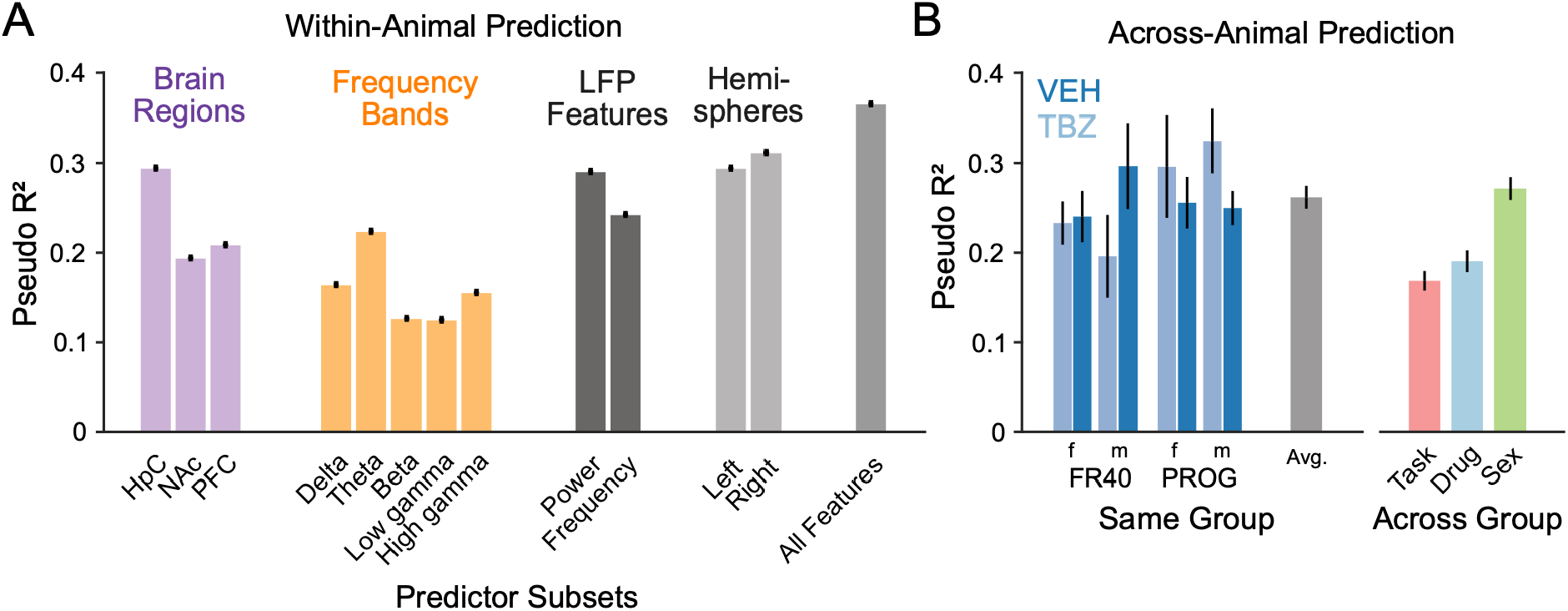
Decoding accuracy. A) Cross-validated within-animal pseudo-R^2^ using all predictors (right) and subsets of predictors from specific brain regions, LFP bands, types (power or peak frequency), and hemispheres. B) Generalization accuracy across animals cross-validated here using leave-one-animal-out (same group) or using a cross-prediction from one group of animals to single animals in another group with one factor modified (across group). Generalization accuracy within the same group does vary slightly between conditions (bottom). Error bars indicate standard error across recording sessions.

In the results above, we used within-animal cross-validation to estimate decoding accuracy. To evaluate the generalization of these models, we next assess accuracy across animals by fitting the full model to data from some animals and testing on a held-out animal. First, we test generalization within groups. That is, we test, for example, how well a model trained on all but one of the male-FR40-VEH recordings performs when tested on the held-out male-FR40-VEH recording (leave-one-animal-out cross-validation). Here we find that there is some variability across groups in generalization accuracy (Fig 4B), but on average, across all groups, the generalization decoding accuracy is relatively high 26.2±1.3% (mean±SE). Additionally, we find that we can generalize across groups that vary in their conditions. Here we train on all of the animals within one group and then assess generalization accuracy using an animal from a group with one factor (task, drug, or sex) modified. That is, we test, for example, how well a model trained on all male-FR40-VEH recordings performs when tested on a held-out male-PROG-VEH recording (across task generalization) or when tested on a held-out female-FR40-VEH recording (across sex generalization). We find that across group generalization is highest when generalizing over sex 27.1±1.3% (mean±SE) and lower, but non-zero, when generalizing over drug (19.0±1.2%) or task (16.8±1.1%).

To understand how the different LFP features are weighted to predict press behavior, we compare the coefficients across single-animal models (Fig 5A). Although there is variability in the coefficients across animals, we find that there is also a substantial conservation of features in these different models. The coefficients reported here act to weigh standardized (z-scored) predictors in the Poisson GLM with an exponential nonlinearity. We estimate group effects using a linear mixed-effects model and report estimated marginal means (see Methods). Increasing hippocampal theta peak frequency by 1 SD corresponds to a decreasing predicted press rate of (−8.7% [-9.7%, -7.6%] left hemisphere and -8.9% [-9.9%, -7.9%] right hemisphere, mean and 95% CI). Increasing hippocampal theta power by 1 SD (−3.3% [-4.5%, -2.2%] left hemisphere and - 4.7% [-5.8%, -3.5%] right hemisphere) and delta power by 1 SD (−2.6% [-3.5%, -1.7%] left hemisphere and -3.9% [-4.8%, -3.0%] right hemisphere) also correspond to decreases in pressing. Across brain regions, increasing low gamma power tends to correspond to decreased pressing, while increasing high gamma power corresponds to increased pressing, but there appear to be some differences in the coefficients for left and right hemispheres and depending on the experimental condition.

**Figure 5.**
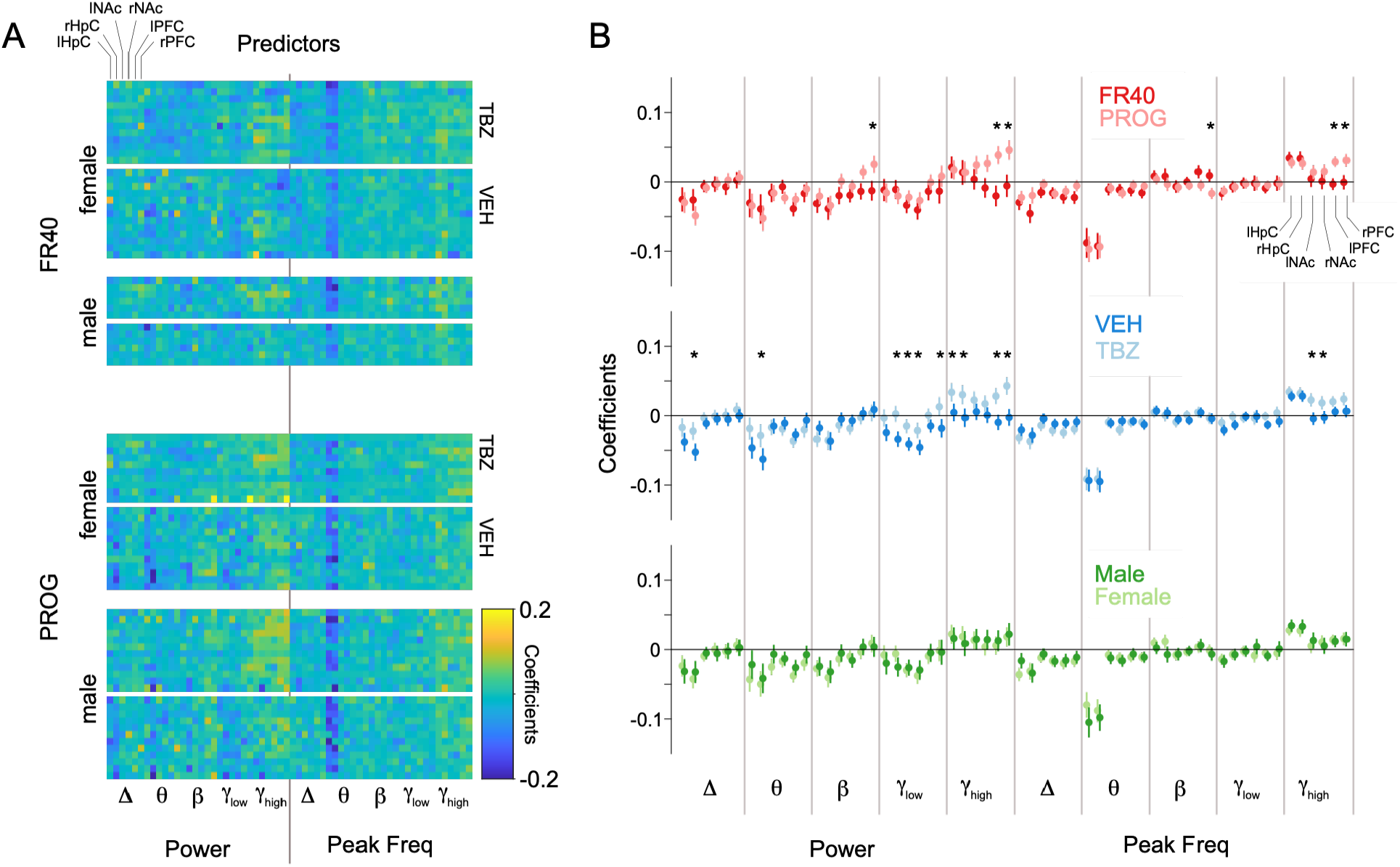
Coefficients across animals. A) Heatmaps illustrate the patterns of standardized coefficients across all recordings in all task, sex, and drug conditions. B) Estimated marginal means for coefficients for task (top), drug (middle), and sex (bottom) comparisons based on linear mixed effects model for each coefficient with random intercepts for each animal. Error bars denote SE. * denotes Bonferroni corrected p<0.05. Coefficients reflect z-scored predictors in the log-linear model.

Comparing coefficients across task, drug, and sex groups, we find that there are some differences, specifically for models that differ in the drug and task (Fig 5B). Comparing FR40 to PROG, we find that the coefficients for right prefrontal cortex show statistically significant task differences in the beta band power (Bonferroni corrected p=0.02) and peak frequency (p=0.02) features. Similarly, we find a statistically significant difference in the high gamma band in both left and right prefrontal cortex for power (p<10^-4^ left and p<0.01 right) and peak frequency (p<10^-4^ left and p<0.01 right). In most of these cases (except beta peak frequency), the coefficient for PROG is greater than the coefficient for FR40. These differences suggest that the association between pressing and the LFP features in the PFC changes somewhat with the task, but these differences are relatively small compared to the overall pattern across all brain regions and features.

Comparing across drug conditions, we find that the coefficients for the right hippocampus are statistically significantly higher under TBZ for both delta power (Bonferroni corrected p<10^-2^) and theta power (p=0.01) compared to VEH. The coefficients for low gamma power are also higher for TBZ compared to VEH for the right hippocampus (p<10^-4^), both left and right nucleus accumbens (p=0.01 left and p=0.03 right) and right prefrontal cortex (p<0.01). Similarly, for the high gamma band power coefficients under TBZ are higher for hippocampus (p=0.045 left and p<0.001 right) and prefrontal cortex (p<10^-7^ for both left and right hemispheres) and peak frequency coefficients are higher in the nucleus accumbens (p<0.001 left and p=0.04 right). These differences suggest that, like the task differences, the association between pressing and the LFP features changes somewhat with the drug. However, the drug differences do not appear to be limited to PFC.

Finally, we compare coefficients across male and female animals. In this case, we do not find any statistically significant differences between groups.

The fact that predictions generalize across task, sex, and drug conditions, suggest that there is some conservation of the LFP features that predict time-varying pressing. However, the differences in the coefficients described above suggest that the LFP features themselves or the association between LFP features and pressing can also shift across conditions. To understand how identifiable the experimental conditions are from their overall coefficients, we fit a logistic model using the coefficient vectors to classify the binary groups in each experimental condition (task, sex and drug). Cross-validated accuracy for task, drug, and sex when classifying based on the coefficients were 86.8%, 77.1%, 60.2% respectively compared to chance levels (with shuffled labels, task=55.4%, drug=53.0%, sex=59.0%). These results illustrate that the coefficient vectors contain some information about the different experimental contexts, but there is still uncertainty about the group membership, especially for male vs female recordings.

One possible explanation for the differences in coefficients across conditions may be that the overall behavior differs across conditions. We thus sought to determine to what extent the coefficient vectors contain information about behavioral summaries in their geometry. For the paired recordings from the same animal under VEH and TBZ, we computed the cosine similarity between coefficient vectors. We find that as the coefficient similarity increases, the degree of behavioral suppression (percent decrease in pressing from VEH to TBZ) decreases (Fig 6A) (correlation coefficient in an arcsine vs Fisher transformed space, r=-0.77, p<10^-8^). This suggests that when TBZ had less of a suppressive impact on lever pressing behavior of an individual animal, the coefficients for the TBZ recording were more similar to those for the VEH recording. Additionally, the geometry of the coefficients themselves appears to partially reflect some individual differences in total pressing (Fig 6B). Here we use multi-dimensional scaling to embed the coefficient vectors in a 1D space. We find that this embedding dimension is correlated with the total pressing (r=-0.76, p=10^-15^ using log-transformed press rate). This trend is partially influenced by the shift in pressing from VEH to TBZ, but even within these drug groups the embedding is correlated with individual pressing (r=-0.40, p<0.01 for VEH and r=-0.73, p<10^-7^ for TBZ).

**Figure 6.**
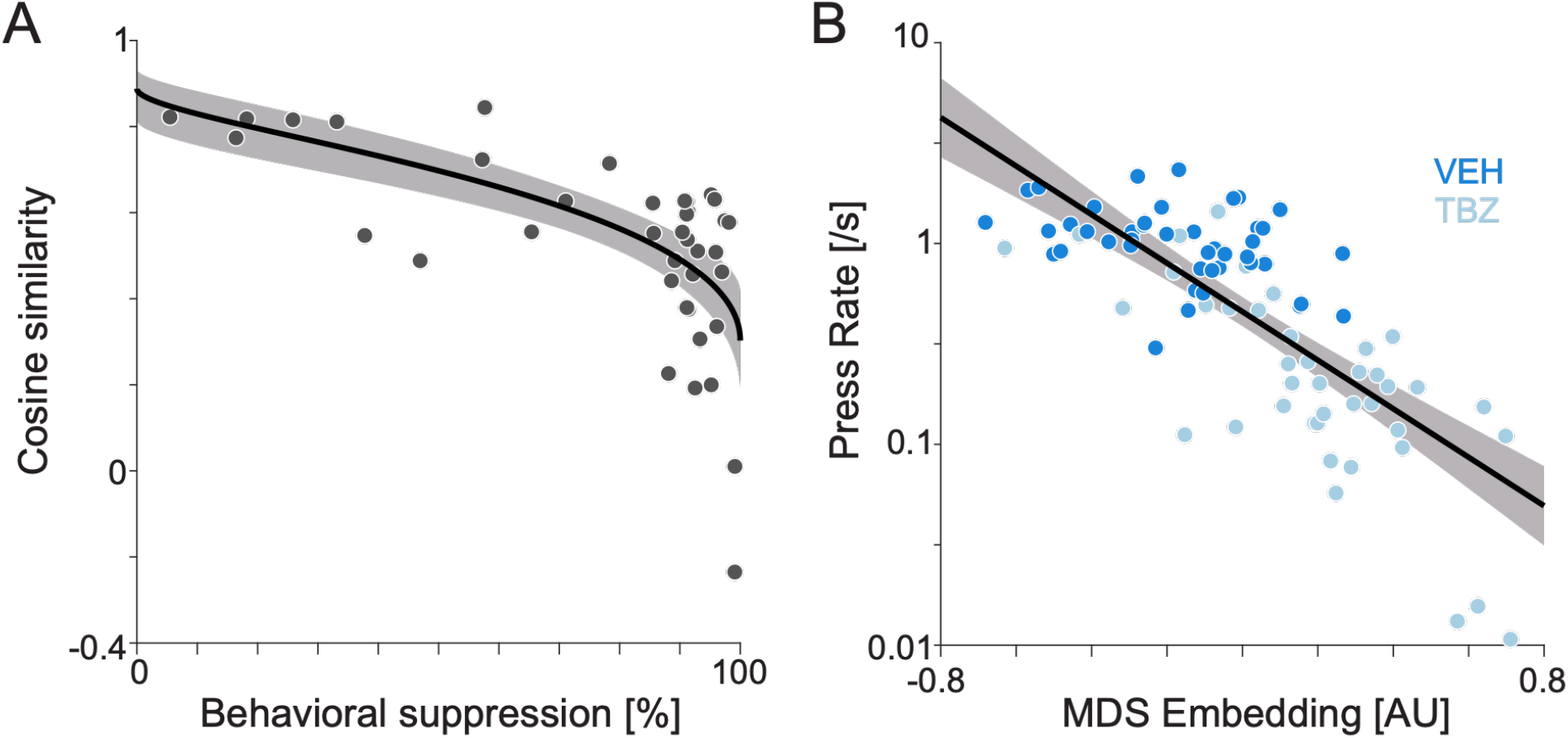
Differences in coefficient vectors reflect individual differences in behavioral summaries. A) Cosine-similarity of the coefficients within the same animal under TBZ vs VEH compared to the extent of behavior suppression under TBZ. To describe the trend, we fit a linear regression in a transformed space (Fisher z-transformed cosine similarity, arcsine transformed behavioral suppression). The curve shows the model fit with the shaded boundary representing standard error. B) 1-dimensional MDS embedding vs press-rate (log scale) for each recording. To describe this trend, we fit a linear regression using log transformed press-rate and the 1-dimensional MDS embedding. Line shows the model fit and confidence band denotes standard error.

One factor that may also affect the results is the timescale of the behavior itself. Animals often alternate between pressing and not pressing, creating autocorrelations in the behavioral data. We, thus, evaluated these autocorrelations and fit delayed models, where the LFP features are shifted in time relative to the observed pressing. We find that the predictions are most accurate in a localized, relatively narrow set of delays between –10s to +10s (Fig 7A). The predictions are, however, slightly longer range than the autocorrelation and, also, tend to have a slight asymmetry where positive delays are predicted more accurately than negative delays. These positive delays correspond to fits where the LFP features at a given time are predicting past pressing.

**Figure 7.**
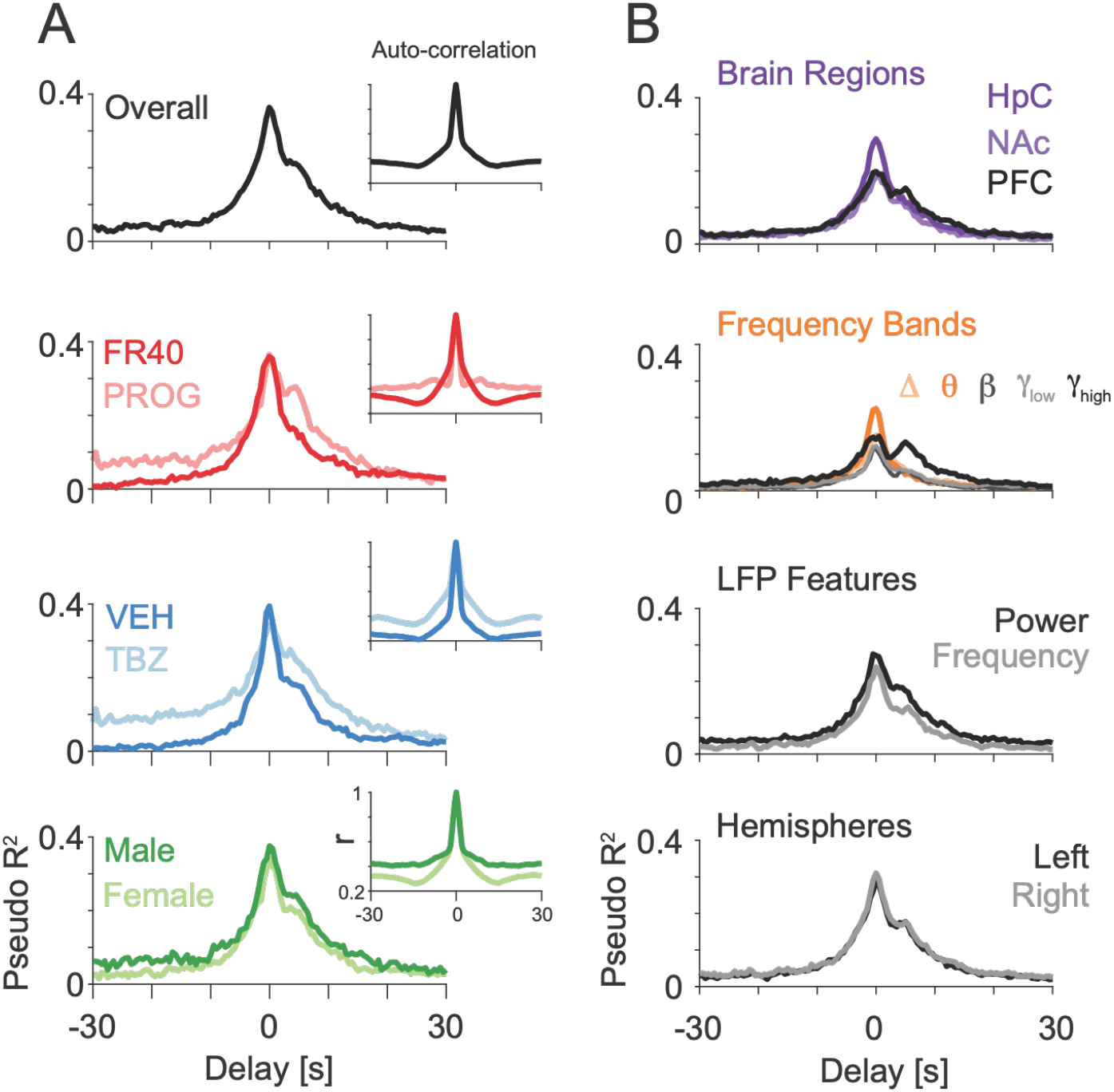
Temporal precision of decoding. A) Median cross-validated pseudo-R^2^ for time-shifted versions of the lever press response variable overall (top) and split across task, drug, and sex conditions. Insets show corresponding auto-correlation for lever pressing. B) Cross-validated pseudo-R^2^ for different delays when modeled using subsets of predictors limited to specific brain areas, frequency bands, LFP features, or brain hemispheres.

We also find that the accuracy of the predictions (pseudo-R^2^) at different delays differs somewhat depending on the experimental condition. Comparing FR40 vs PROG, we find that PROG has higher prediction accuracy across all the delays while having similar accuracy to FR40 at zero delay. Comparing VEH vs TBZ, we also find TBZ having higher prediction accuracies at all delays while matching VEH accuracy at zero delay. For male vs female comparison, we find that male shows higher accuracy values throughout positive and negative delays while matching with female accuracy at zero delay. These trends mostly mirror the difference in autocorrelation between groups, where TBZ and PROG recordings have longer timescale correlations compared to VEH and FR40 recordings.

As in the predictions at zero delay, described above, the delayed predictions differ when we restrict the predictors to specific brain regions, frequency bands, or features (Fig 7B). Across brain regions, although at 0 delay the best prediction accuracy comes from using LFP features form the hippocampus, at a delay of 5s the PFC has the highest accuracy. Similarly for frequency bands, although the theta band had the highest prediction accuracy at zero delay, the high gamma band has the highest prediction accuracy at 5 sec delays. Comparing power and frequency features, we find that using power features gives slightly higher accuracy than peak frequency at all delays, but the differences across hemispheres at different delays are minimal.

In addition to predicting press rate directly, it is possible that LFP predictors may contain information about longer timescale representations of the lever pressing behavior, potentially related to the timing of past or future reinforcement or to accumulated effort. To test this possibility, here, rather than decoding press rate, we modified the decoder to predict the 1) cumulative presses since the previous reinforcement delivery or 2) presses remaining until the next reinforcement delivery. These signals differ from the instantaneous press rate, and, also, differ substantially depending on the task (Fig 8A-B). For FR40, the long-term task variables for presses since previous and presses until next reinforcer follow a consistent trajectory that mirrors the animals progress through the ratio. For PROG, as the ratios require progressive more presses, both of the long-term task variables will increase, as well.

**Figure 8.**
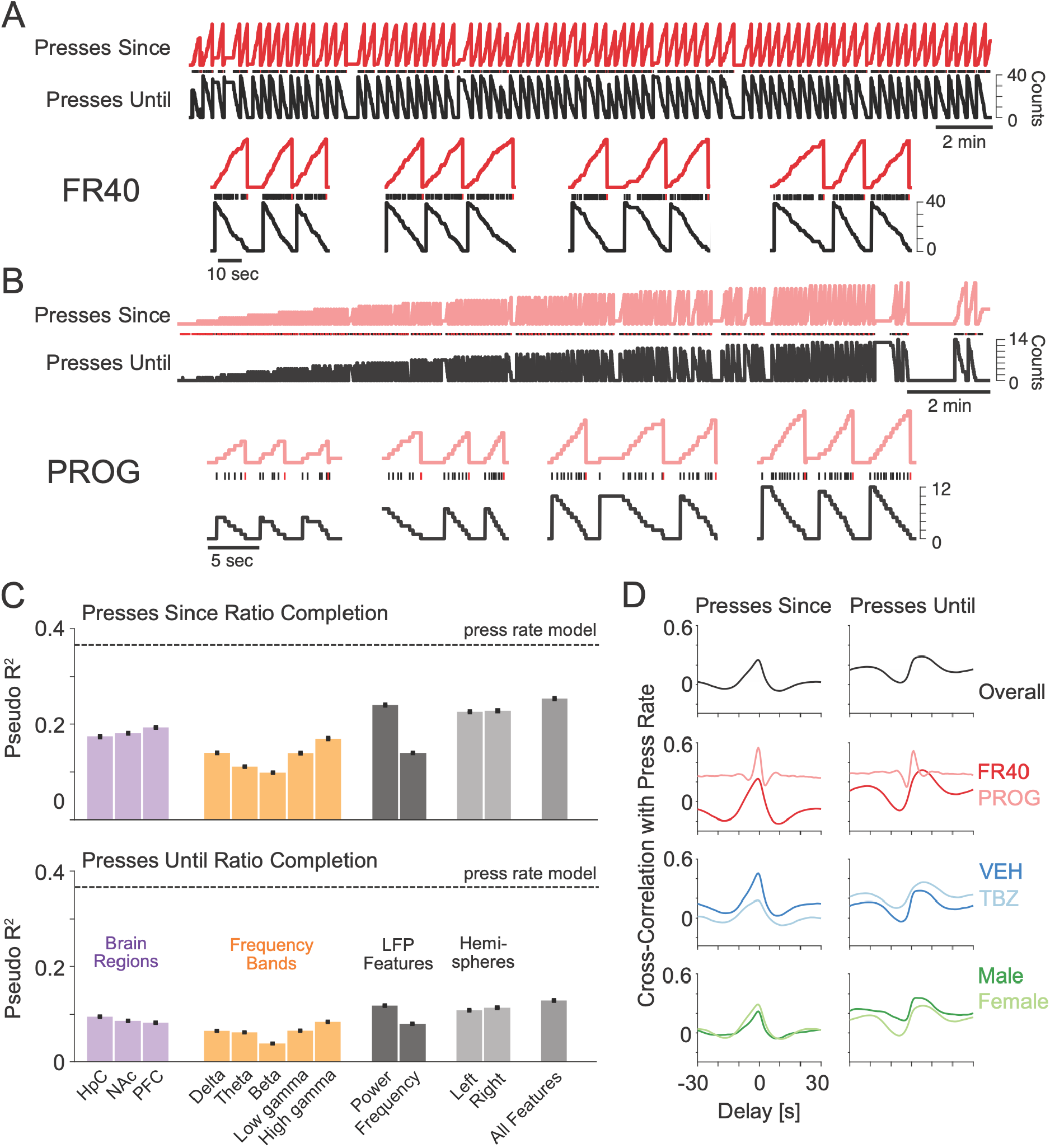
Decoding long-term task representations. A) Example traces for two calculated variables: cumulative press counts since the last reinforcer (red) and presses until the next reinforcer (black) for an example male animal performing the FR40 task. B) Example recording from a female animal performing the PROG task. Both examples show VEH data with the full sequence of the variable (top) and highlighted examples of three complete ratios (bottom). Black ticks denote individual lever presses and red ticks denote ticks that complete the ratio. C) Mean cross-validated pseudo-R^2^ for decoding the presses since ratio completion (top) for subsets of predictors and presses until ratio completion (bottom). Dashed lines denote the pseudo-R^2^ for decoding press rate with all predictors. Error bars denote standard error across recording sessions. D) Cross-correlation between the calculated variables and the press rate for presses since (left) and presses until (right) reinforcement. Results are shown for all recordings (top) and for specific task, drug, and sex conditions.

Here we find that we can decode both calculated long-term task variables. Predictions are not as accurate as predictions of the press rate, but presses since the last reinforcement can be decoded with pseudo-R^2^=25.4±0.2 using all predictors. Presses until the next reinforcement can be decoded with pseudo-R^2^=12.9±0.1 using all predictors. Additionally, as with decoding press rate, the performance varies when the set of predictors is restricted (Fig 8C). Interestingly, the accuracy for presses since the previous reinforcer is highest when LFP features are limited to PFC, unlike decoding for the press rate itself, which was highest for HpC. Accuracy is also higher when decoding using the high gamma band for presses since last reinforcer and presses until next reinforcer, rather than the theta band, which was highest for press rate itself.

These results suggest that longer-term task variables can be decoded from these multi-region LFP features but may rely on a somewhat distinct set of predictors, since we find that using different predictor combinations within brain regions or frequency bands and features lead to different pseudo-R^2^ patterns compared to the press rate models. To quantify the differences between the coefficients we calculate the cosine similarity between the coefficient vectors for the decoders for long-term task variables and press rate itself within the same recording. For presses since the last reinforcer the average similarity is 0.24±0.05, and for presses until the next reinforcer the average similarity 0.26±0.03. These similarities indicate that the coefficients for the long-term task variables somewhat resemble the coefficients for the press rate itself. However, the coefficients for the two long-term variables are dissimilar from each other (average cosine similarity 0.03±0.01), and the similarities of the long-term variable decoders to the press rate decoders are less than those of the press rate decoder coefficients from different recordings (0.32±0.01).

Although these results suggest that LFP predictors may contain information about longer term task representations, it is important to note that these calculated variables are correlated with the press rate itself (Fig 8E). As with the autocorrelations in press rate, there are substantial cross-correlations between the accumulated pressing and the press rate and these correlations differ across conditions. PROG shows consistently higher cross-correlation between long-term variables and the press rate compared to FR40, and TBZ shows higher cross-correlation compared to VEH. The cross-correlation for presses until the next reinforcer peaks at ∼5s delays and also has lower cross-correlation at 0 delay compared to presses since the previous reinforcer. These differences may partially explain why the differences in decoding accuracy for the long-term task variables, and the presence of the correlations may be one reason long-term task variables can be decoded at all.

## Discussion

Here we find that LFP power and peak frequency in the HpC, PFC, and NAc can be used to accurately decode lever pressing behavior during operant performance. The features used for decoding are largely conserved across animals with small differences across task and drug conditions and no detectable differences between male and female rats. We find that across animals, even when the task (FR40 vs PROG) or drug condition (VEH vs TBZ) is varied, decoders can generalize to predict moment-by-moment lever pressing. We find that the best decoding performance is achieved using LFP features from multiple brain areas and multiple frequency bands, including both power and peak frequency features. Additionally, the geometry of the coefficients across animals appears to reflect individual differences in overall behavior and individual differences in the magnitude of drug effects. Overall, these results demonstrate that a distributed, conserved pattern of LFP activity is associated with lever-pressing rate, and this pattern is robust across different animals and contexts.

Decoders based on LFP activity in motor cortex have been used to predict reaching movement kinematics (Sanes and Donoghue, 1993; Mehring et al., 2003; Rickert et al., 2005; Flint et al., 2012). Decoders based on LFPs in humans have been used to predict intended speech (Wilson et al., 2020), and decoders based on hippocampal LFP were reported to be related to spatial position (Agarwal et al., 2014) across individual animals. An important aspect of the decoders described above is their ability to generalize across animals and task, sex, and drug conditions. In many domains, decoders based on single neuron spiking often perform extremely well on individual animals (Glaser et al., 2020), but generalizing decoders across animals is challenging due to the heterogeneity of single neuron responses. Previous work has found that spike-based decoders can generalize when aligning to a common low-dimensional structure (Chen et al., 2021; Safaie et al., 2023). In the present paper, we find that coarse LFP patterns distributed across multiple brain regions have some conserved structure during lever pressing even without additional processing.

Many previous studies have found that hippocampal theta power and peak frequency are positively associated with movement variables (Vanderwolf, 1969b), and report a positive correlation between these features and locomotor speed, in particular. Our present results suggest a negative association for lever pressing, in which increases in theta power and frequency correspond to decreases in local press rates. Previous research reported that increased theta power at the onset of lever pressing was followed by a decrease in rats performing a visual discrimination task (Wyble et al., 2004). Increases in NAc and PFC theta power have also been observed during lever pressing and anticipation of responding (Gruber et al., 2009), and are modulated by wait period and outcome in animals performing a reaction time task (Donnelly et al., 2014). These studies have also found that changes in power occur for other LFP bands, including delta and beta, as does coherence across regions. Interestingly, the present results showed that moment-by-moment lever pressing is best predicted by a multi-region, multi-band pattern rather than any specific region or band in isolation.

Oscillatory activity in the hippocampus (Lopes Da Silva and Kamp, 1969) and, particularly, in the PFC and NAc, has also been associated with reward prediction and goal-seeking (Goto and Grace, 2008; Gruber et al., 2009). In addition to the accurate decoding of immediate lever pressing, we find weaker, but still reliable, LFP patterns associated with longer timescale task variables, such as presses since the last reinforcer and presses until the next reinforcer. The longer-term task variables analyzed here could potentially be associated with changes in exertion of effort and motivational states. However, we do not explicitly alter reward predictability or delivery during the task, and the strong correlation between the long-term variables and immediate lever pressing makes it difficult to determine if there is a unique representation of effort valuation or aspects of motivation. Nonetheless, in view of the well characterized ability of TBZ to deplete dopamine by blocking vesicular storage (Nunes et al., 2013), the differences in the press rate decoder coefficients that we observed between VEH and TBZ conditions may reflect a broader reorganization that mirrors drug-induced changes in dopamine transmission (Salamone et al., 2007). Moreover, task-related differences in press rate decoder coefficients may reflect the distinct lever pressing requirements of the FR40 vs. the PROG operant schedules. Although both schedules require high levels of lever pressing output for obtaining reinforcement, the FR schedule maintains a constant requirement, while the PROG schedule involves increments in the ratio requirement across the session.

Although we have demonstrated that lever pressing can be decoded across a wide range of recording conditions, there are several limitations for the results here. First, even though we remove strong electrical artifacts from our data, the LFP may still be partially influenced by movement and muscle artifacts, including artifacts related to consumption of the reinforcer pellet. However, the periods when the animal is moving around the operant chamber or actively consuming the pellet are small compared to the overall recording period, and lever pressing can be decoded from LFP features even during other task periods and even when limited to low-frequency features only. Second, although we focus on band power and peak frequency, other LFP features such as functional connectivity and cross-frequency coupling (Canolty and Knight, 2010; Michaels et al., 2018) could potentially improve decoding beyond what we show here for band power and peak frequency. Recent work has also suggested that taking aperiodic activity into account (Donoghue et al., 2020) may provide more robust estimates of these within-band LFP features. Finally, the present work focused on three brain regions (HpC, PFC, and NAc), but other areas involved in operant behavior and motivation, such as the amygdala (Baxter and Murray, 2002; Janak and Tye, 2015) and neostriatum (Balleine et al., 2007), likely also contain task-associated changes in LFP (Berke, 2009; Leventhal et al., 2012).

Identifying robust, functionally meaningful multi-area patterns of neural oscillations is a major, ongoing goal of systems neuroscience (Buzsáki and Draguhn, 2004). In humans, unique multi-region patterns have been identified associated with resting states (Fox et al., 2005; Raichle et al., 2001) and attention (Vossel et al., 2014) among others. The results here are particularly relevant, to potential networks and biomarkers related to depression and Parkinson’s disease. Dopamine depletion during operant behavior has been previously used as a model of depressive symptoms related to motivational dysfunction (Salamone and Correa, 2024). Although here we find a clear pattern of conserved LFP activity across the conditions measured here, there are also some consistent variations due to task (FR40 vs PROG) and due to TBZ, as well as individual differences across animals. Since these conditions contain alterations in motivation, it may be worthwhile to examine whether comparable changes in oscillatory activity occur in human depression.

## Acknowledgments

Research reported in this publication was supported by the National Institute of Mental Health of the National Institutes of Health under award number 5R01MH121350. The content is solely the responsibility of the authors and does not necessarily represent the official views of the National Institutes of Health. Research was additionally supported by grants from the University of Connecticut Research Foundation and the Connecticut Institute for Brain and Cognitive Sciences.

