## Supplementary Figures for "Decoding instrumental lever pressing from prefrontal, accumbens, and hippocampal local field potentials: conservation across sex, task, and dopamine depletion"

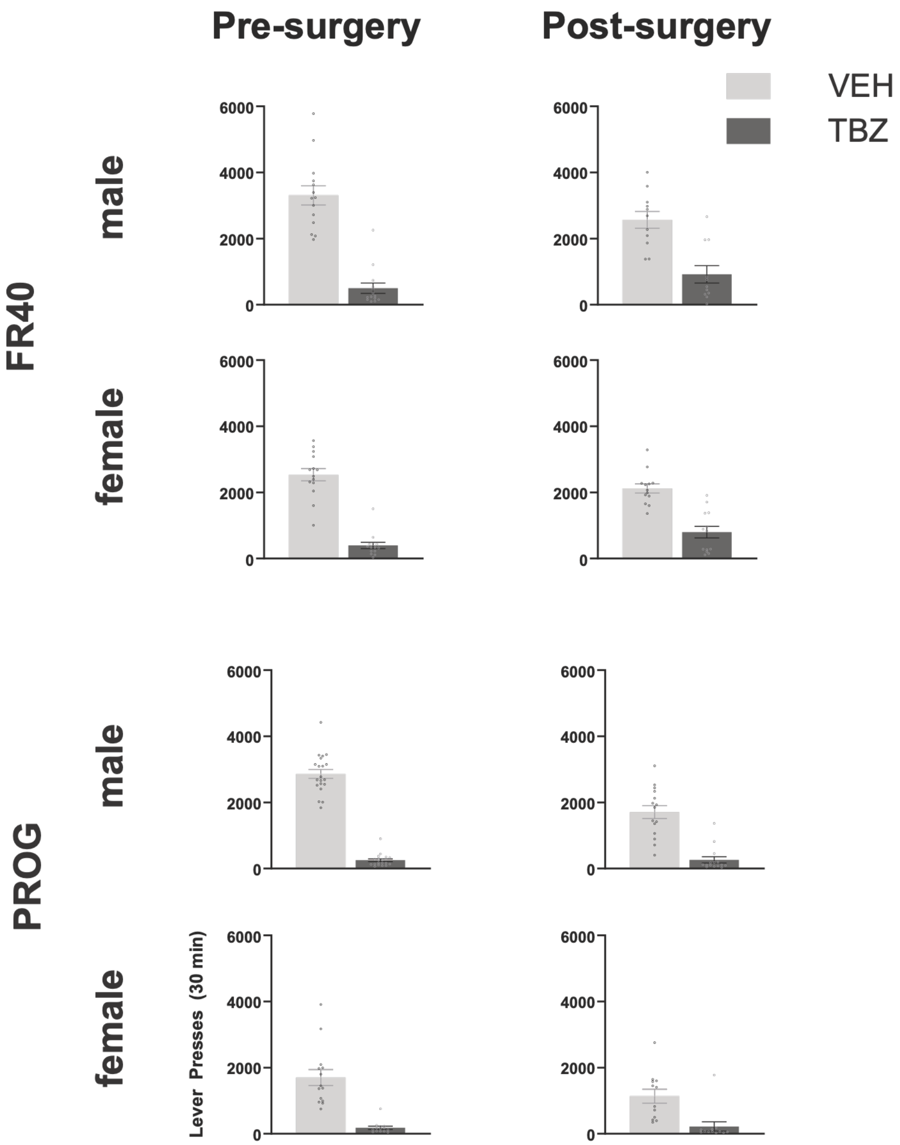


**Figure S1.** Lever pressing behavior pre- and post-surgery. Number of lever presses during a 30-minute recording pre-surgery (left) and post-surgery (right) sessions. Error bars indicate standard error. TBZ and VEH differences are all statistically significant (paired t-test, $p<0.05$).

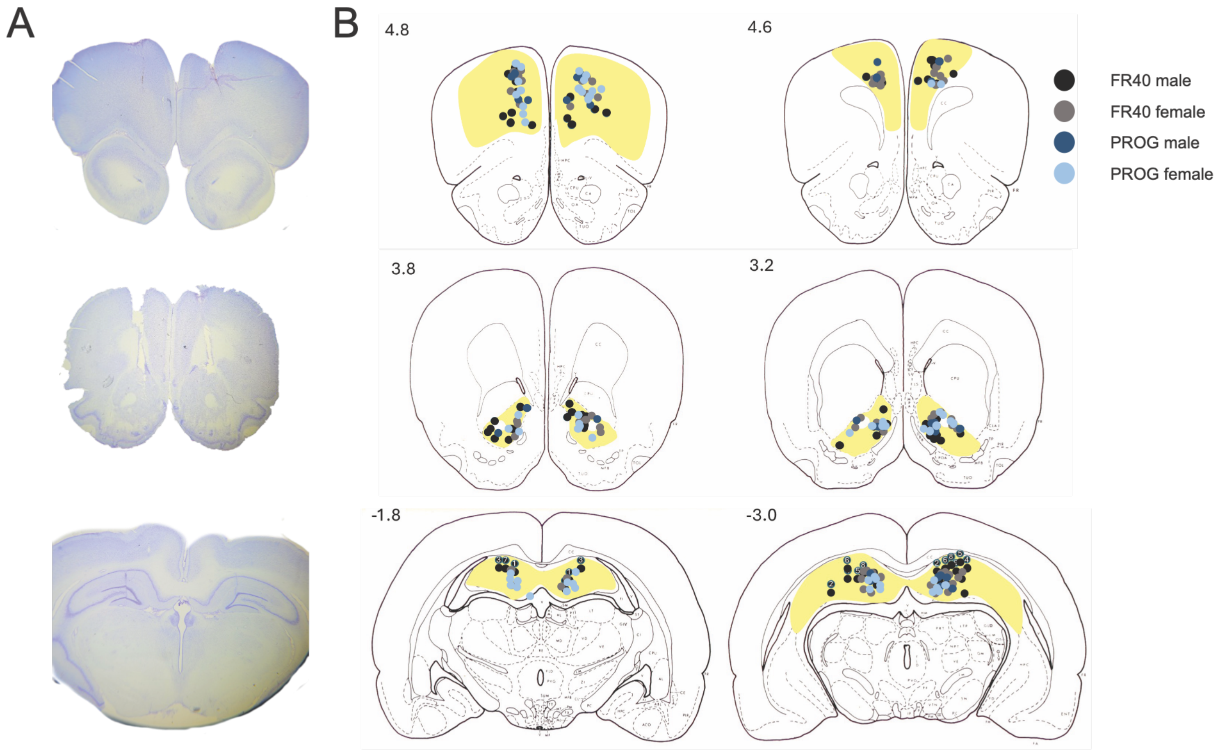


**Figure S2.** Histology. A) Nissl-stained slices showing electrode tracts in medial Prefrontal Cortex (top), Nucleus Accumbens (middle) and dorsal Hippocampus (bottom) from different exemplar animals. B) Distribution of electrode placements in medial Prefrontal Cortex (top), Nucleus Accumbens (middle) and dorsal Hippocampus (bottom) for the animals in each task and sex condition.
